# Host obesity impacts genetic variation in influenza A viral populations

**DOI:** 10.1101/2023.07.12.548715

**Authors:** Marissa Knoll, Rebekah Honce, Victoria Meliopoulos, Stacey Schultz-Cherry, Elodie Ghedin, David Gresham

## Abstract

Obesity is a chronic health condition characterized by excess adiposity leading to a systemic increase in inflammation and dysregulation of metabolic hormones and immune cell populations. Obesity is well established as a risk factor for many noncommunicable diseases; however, its consequences for infectious disease are poorly understood. Influenza A virus (IAV) is a highly infectious pathogen responsible for seasonal and pandemic influenza. Host risk factors, including compromised immunity and pre-existing health conditions, can contribute to increased infection susceptibility and disease severity. During viral replication in a host, the negative sense single stranded RNA genome of IAV accumulates genetic diversity that may have important consequences for viral evolution and transmission. Here, we investigated the impact of host obesity on IAV genetic variation using a diet induced obesity ferret model. We infected obese and lean male ferrets with the A/Hong Kong/1073/1999 (H9N2) IAV strain. Using a co-caging study design, we investigated the maintenance, generation, and transmission of intrahost IAV genetic variation by sequencing viral genomic RNA obtained from nasal wash samples over multiple days of infection. We found evidence for an enhanced role of positive selection acting on *de novo* mutations in obese hosts that led to nonsynonymous changes that rose to high frequency. In addition, we identified numerous cases of recurrent low-frequency mutations throughout the genome that were specific to obese hosts. Despite these obese-specific variants, overall viral genetic diversity did not differ significantly between obese and lean hosts. This is likely due to the high supply rate of *de novo* variation and common evolutionary adaptations to the ferret host regardless of obesity status, which we show are mediated by variation in the hemagglutinin (HA) and polymerase genes (PB2 and PB1). As with single nucleotide variants, we identified a class of defective viral genomes (DVGs) that were found uniquely in either obese or lean hosts, but overall DVG diversity and dynamics did not differ between the two groups. Our study provides the first insight into the consequences of host obesity on viral genetic diversity and adaptation, suggesting that host factors associated with obesity alter the selective environment experienced by a viral population, thereby impacting the spectrum of genetic variation.

## Introduction

Obesity is an increasing public health concern as the number of people who are considered obese has nearly tripled since 1975, with over 600 million adults considered overweight in 2015 (The GBD 2015 Obesity Collaborators, 2017). Although primary health risks associated with obesity include cardiac and metabolic syndromes, obesity is also known to alter immune function through the endocrine action of adipocytes, which are both enlarged and more numerous in obesity (Martí et al., 2001; Rajala & Scherer, 2003). Adipocytes release immune related proteins, including leptin, which has been shown to upregulate the production of pro-inflammatory cytokines (Loffreda et al., 1998; Lord et al., 1998). In addition, adipocytes secrete higher levels of immune proteins both locally, including TNF-α, and systemically, including C-reactive protein, interleukin 6 (IL-6), and serum amyloid A (SAA) (Rajala & Scherer, 2003; Hotamisligil & Spiegelman, 1994). Obesity is also linked to dysregulation of immune cells, such as reduced lymphocyte proliferation and elevated leukocyte and lymphocyte counts (Hotamisligil & Spiegelman, 1994; Nieman et al., 1999), putatively due to the increased systemic inflammation and dysregulated metabolic hormones. Obesity has been shown to exacerbate noncommunicable disease, such as cardiovascular disease, metabolic syndromes, cancer, as well as infectious diseases (McClean et al., 2008; Gonzalez, 2006; McClean et al., 2008; Ogden et al., 2007). The link between host obesity and infectious diseases is especially interesting as changes in immune function could contribute significantly to infection dynamics and disease severity.

Influenza A virus (IAV), a single stranded, negative sense segmented RNA virus, is a common respiratory pathogen that causes infection of the upper and, in severe cases, the lower respiratory tracts (Paules & Subbarao, 2017). Obesity is implicated in more severe disease and increased mortality in IAV-infected individuals (Liu et al., 2014; Morgan et al., 2010; Louie et al., 2011; Van Kerkhove et al., 2011). However the underlying mechanisms have not been fully elucidated. Case studies and population data have shown increased IAV shedding, indicative of higher viral loads, in obese patients (Fleury et al., 2009; Maier et al., 2018). In addition, obese patients have lower type I interferon production and delayed antiviral responses (Terán-Cabanillas et al., 2014). These factors may impair the ability of an obese host to suppress viral replication, resulting in delays in viral clearance and increases in the length of illness. Animal studies provide additional evidence for these effects as high calorie diets led to altered metabolic and immunological profiles and increased mortality in response to IAV infection (Hariri & Thibault, 2010; Lumeng et al., 2007; Smith et al., 2007).

As with all RNA viruses, the error prone nature of the RNA-dependent RNA polymerase in IAV can lead to significant within-host genetic diversity of the viral population (Webster et al., 1992). This genetic diversity can take many forms including single nucleotide variants (SNVs) and large internal deletions known as defective viral genomes (DVGs). The genetically diverse population within a host experiences selective pressure from the host environment, leading to allele frequency changes of SNVs, especially those in the genes that code for the antigenically variable surface proteins hemagglutinin (HA) and neuraminidase (NA) (Webster et al., 1992). By contrast, DVGs are predominantly found in the polymerase gene segments (PB2, PB1, PA). The extent to which the altered metabolic and immune status of obese hosts impacts the accumulation and selection of IAV genetic variation has not previously been investigated. The altered immune state in the obese host may result in a distinct environment compared to the lean host that could alter the dynamics and diversity of genetic variation through differential selection.

To study the accumulation of genetic variation in IAV populations over the course of infection, we developed a diet-induced obesity model in ferrets (Meliopoulos et al., *unpublished*). The use of ferrets as a model for human IAV infections is well established and has a number of advantages, including their susceptibility to and transmissibility of human-relevant strains without extensive initial host adaptation (Huber & McCullers, 2006; Kirchenbaum et al., 2017; Paquette et al., 2014; Peng et al., 2014; Kirchenbaum & Ross, 2017). Thus, using an obese ferret model allows us to examine IAV infections in a physiologically-relevant context for human obesity.

Human IAV infections are most frequently caused by H1N1 and H3N2 subtypes; however, many other subtypes can cause disease and thus are of major health concern. H9N2 is a low-pathogenicity avian-like influenza virus with reservoirs in wild bird populations and domestic poultry, which has led to occasional human infections (Carnaccini & Perez, 2020). Although there is no current evidence for human-to-human transmission of H9N2, it remains a virus of pandemic concern by the CDC (*Viruses of Special Concern*, 2019). Ferret models have been used to investigate H9N2 infectivity and transmissibility (Wan et al., 2008). However, the effect of obesity on the genetic diversity in the viral population during H9N2 infection has not previously been studied.

To investigate the effect of host obesity on genetic diversity, we infected neutered male obese ferrets and lean ferrets with the A/Hong Kong/1073/1999 (H9N2) strain. Using a co-caging study design, we investigated the generation and transmission of intrahost genetic diversity in obese and lean animals. We assayed genetic diversity in the viral population through optimized experimental and computational workflows to quantify intrahost SNV (Roder et al., 2023) and DVG (https://github.com/GhedinSGS/DiVRGE) diversity from Illumina sequencing data across multiple days of infection and transmission pairs. We found evidence of strong positive selection in the form of nonsynonymous SNVs that increased to high frequencies in obese contact ferrets independent of transmission, and multiple recurrent low frequency variants that were unique to obese ferrets. In addition, we demonstrated that adaptation to the ferret host was mediated by variation in HA and polymerase genes (PB2 and PB1) that was independent of diet. Despite these differences, SNV and DVG diversity did not differ in obese hosts compared with lean hosts. Our study suggests that host factors associated with obesity alter the selective environment experienced by the viral population, resulting in a unique class of obese-specific genetic variants.

## Results

### Study Design

We investigated the effect of host obesity on IAV genetic diversity during infection using a ferret obesity model. Briefly, neutered male ferrets were randomly assigned to the lean or obese group and reared on the appropriate diet for 12 weeks (Karlsson et al., 2014) (**Fig.1A**). Infection experiments were performed in 5 cohorts over the course of roughly 2.5 years (Fall 2017, Winter 2017, Summer 2018, Spring 2019, and Spring 2020) (**Table S1**). To generate a stock of virus, the H9N2 strain A/Hong Kong/1073/1999 was propagated in 9-day old embryonated chicken eggs and then stored. One aliquot per cohort was thawed and diluted in PBS for use as the inoculum in our experiments. Index ferrets were infected intranasally with 10^6^ tissue culture infectious dose-50 (TCID_50_) of the A/Hong Kong/1073/1999 (H9N2) inoculum. On the first day post-infection (dpi), an influenza-naive contact ferret was introduced to the same cage as a single index ferret. Our experimental design included all pairwise combinations of index and contact ferrets; namely, lean index and lean contact, lean index and obese contact, obese index and lean contact, and obese index and obese contact (**Table S2**). Index and contact ferrets remained co-caged until the end of the experiment at 12 dpi. Viral RNA was extracted from nasal wash samples collected every second day from 2 dpi to 12 dpi from both index and contact ferrets. All RNA samples were reverse transcribed, PCR amplified, and sequenced in duplicate. RNA from the viral inoculum was also extracted and sequenced to define pre-existing diversity.

**Figure 1.**
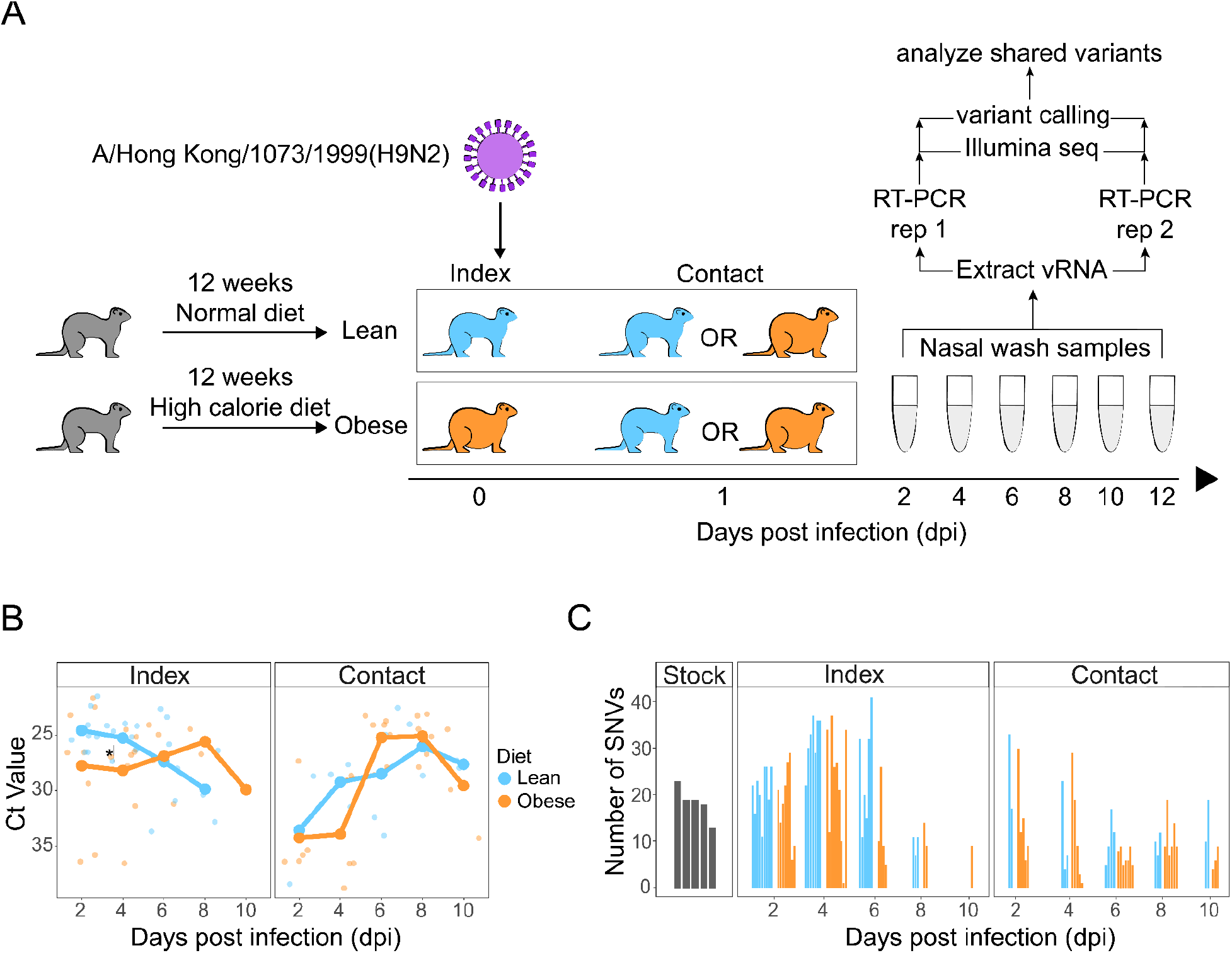
Infection of lean and obese ferrets with A/Hong Kong/1073/1999 (H9N2) virus. **A)** Schema of experimental design: Influenza-naive neutered male young adult ferrets were fed a lean or obese diet for 12 weeks prior to infection with A/Hong Kong/1073/1999 (H9N2) influenza virus. Index ferrets were inoculated intranasally with 10^6^ TCID_50_ on day 0. An index ferret was co-caged with a contact ferret at 1 dpi. All possible combinations of lean and obese index and contact ferrets were tested. At two day increments from 2 - 12 dpi, nasal wash samples were collected, viral RNA was isolated, reverse transcribed, amplified, and sequenced in duplicate. **B)** Viral load was quantified using RT-PCR analysis of the M gene and estimated using the critical threshold (Ct) method for all samples passing sequencing quality thresholds. Small points indicate values for individual samples. Large points connected by lines are average values of samples for each diet group (obese and lean). Facets indicate infection route and color indicates diet. * = p < 0.05 using a two-tailed Student’s t test. **C)** Number of SNVs identified per sample (n = 81) at each time point. The five different stock values correspond to the five inocula for the different cohorts used in our study. Individual samples from ferret nasal washes are sorted by dpi.

We studied a total of 28 index (15 obese and 13 lean) and 20 contact (12 obese and 8 lean) ferrets over the course of the experiment acquiring a total of 180 nasal wash samples from 48 ferrets over 2 - 12 dpi. Each ferret was assigned a unique identification code (**Table S1**). After applying sequencing quality thresholds (see **Methods**), our dataset comprised 37 ferrets for which at least one sample was suitable for SNV analysis resulting in a total of 81 successfully sequenced samples. In general, failure to acquire sequence data was due to either unsuccessful infection of the index ferret, unsuccessful transmission, or clearance of the infection in infected ferrets at later time points in the experiment. All sequenced samples were acquired from 2-10 dpi and no sequencing data was successfully obtained from samples at 12 dpi suggesting that the infection had been cleared in all individuals by this point. Our sequencing dataset includes 11 successful transmission pairs, for which both index and contact ferrets had at least one successfully sequenced sample (**Table 1**). The average sequence coverage across all 8 genome segments exceeded 1,000-fold enabling sensitive detection of low frequency variants (**Supp. Fig. 1A**).

**Table 1.** Distribution of the 37 unique ferrets with successful IAV sequencing. The number of obese and lean index and contact ferrets for which high quality sequencing data was acquired. A successful transmission event is defined by the acquisition of sequence data for both an index ferret and its transmission partner. In some cases data was obtained for a contact ferret but not for the corresponding index ferret and therefore was not considered a positive transmission event.

| Index Diet | Transmission Events | Contact Diet |
| --- | --- | --- |
| Obese<br>(n = 10) | Obese to obese<br>(n = 7) | Obese<br>(n = 7) |
| Obese<br>(n = 2) | Obese to lean<br>(n = 0) | Lean<br>(n = 1) |
| Lean<br>(n = 4) | Lean to obese<br>(n = 2) | Obese<br>(n = 3) |
| Lean<br>(n = 7) | Lean to lean<br>(n = 2) | Lean<br>(n = 3) |

To quantify viral load we performed quantitative RT-PCR of the M genomic segment and determined the critical threshold (Ct) value for all 81 samples that passed quality control filters (**Fig. 1B**). Lower Ct values indicate more viral RNA material. We found that the overall trend in Ct does not differ significantly between obese and lean index and contact ferrets with the exception of 4 dpi in the index ferrets. This suggests that total infectious viral load did not differ between lean and obese hosts.

We performed single nucleotide variant detection on all 81 samples using a 1% allele frequency cutoff, requiring that variants were present in both technical replicates (Roder et al., 2023). Inoculum samples used to infect index ferrets are expected to be genetically heterogeneous as a result of amplification in cell culture. Indeed, we identified an average of 18.4 SNVs in the 5 different inoculum aliquots (**Fig. 1B**). Variant calling for all ferret samples was performed relative to the consensus sequence of the stock for each cohort. Samples contained an average of 18 SNVs (**Fig. 1C**). As expected, viral Ct values were negatively correlated with the number of SNVs in index ferrets, as a larger population size (i.e. more RNA molecules resulting in a lower Ct value) is expected to correspond to more genetic diversity (**Supp. Fig. 1B**). Interestingly, this was not the case for contact ferrets, as Ct values were not correlated with the number of variants. When comparing similar viral abundances, we observe systematically reduced diversity in contact ferrets compared with index ferrets possibly reflecting differences related to direct and transmitted infection (**Supp. Fig. 1B**).

### Obesity selects for unique consensus changes

As obesity has been shown to lead to significant differences in gene expression and immune function, we hypothesized that different host selection pressure in obese ferrets may result in diet-specific adaptation. Therefore, we identified SNVs that were unique to either the obese or lean diet groups. We distinguish between consensus changes, in which 50% or more of sequence reads at a specific site contain a different nucleotide than the viral inoculum, and minor variants, in which fewer than 50% of sequence reads at a specific site contain a different nucleotide than the viral stock.

We first investigated consensus changes as the high frequency of these SNVs could be a result of positive selection. There were 13 consensus changes identified in 8 of the 37 ferrets (9 in obese ferrets and 4 in lean ferrets) (**Table 2**). Among these 8 ferrets, 4 of them had more than one consensus change. Surprisingly, the vast majority (12 of 13) of consensus changes were found in contact ferrets. These appear to be *de novo*, and not transmitted, variants as they were not present in their index transmission partner at any frequency. Furthermore, two of these variants occurred recurrently in 2 independent ferrets. One of these, L628M, is a nonsynonymous mutation that reached high frequency (> 85%) in both ferrets, whereas the other, F9F, is a synonymous mutation that was present at lower frequency in both ferrets (< 67%).

**Table 2.** Sequence consensus changes identified in infected ferrets. The 13 consensus changes (allele frequency greater than or equal to 50%) at the nucleotide and amino acid level found in 8 ferrets. Black borders delineate individual ferrets. Recurrent consensus changes are indicated in bold.

| Segment | Mutation Type | Nucleotide Change | Amino Acid Change | Frequency | Infection Route | Diet | Ferret ID |
| --- | --- | --- | --- | --- | --- | --- | --- |
| **PB1** | **Nonsynonymous** | **C1882A** | **L628M** | **92%** | **Contact** | **Obese** | **1409** |
| NP | Nonsynonymous | C326T | T109I | 86% | Contact | Obese | 1409 |
| HA | Nonsynonymous | C284A | S95Y | 52% | Contact | Obese | 1794 |
| PB2 | Nonsynonymous | C334T | P112S | 99% | Contact | Lean | 1797 |
| PB1 | Nonsynonymous | T965C | I322T | 75% | Contact | Obese | 1980 |
| PA | Synonymous | A1483C | R495R | 80% | Contact | Lean | 1986 |
| NS | Nonsynonymous | C652T | Q218X | 83% | Contact | Lean | 1986 |
| **PA** | **Synonymous** | **C27T** | **F9F** | **67%** | **Contact** | **Obese** | **2231** |
| NS | Synonymous | C267T | Y89Y | 62% | Contact | Obese | 2231 |
| PB2 | Synonymous | C639A | R213R | 61% | Contact | Obese | 2232 |
| **PB1** | **Nonsynonymous** | **C1882A** | **L628M** | **85%** | **Contact** | **Obese** | **2232** |
| HA | Synonymous | A747G | Q249Q | 56% | Contact | Obese | 2232 |
| **PA** | **Synonymous** | **C27T** | **F9F** | **51%** | **Index** | **Lean** | **2254** |

About half (7 of 13) of the consensus changes were nonsynonymous and were located in the PB2, PB1, HA, NP, and NS segments (**Fig. 2A**). Nonsynonymous consensus changes tended to be found at higher allele frequencies than synonymous consensus changes (**Fig. 2B**). Three of the consensus changes showed an increase in allele frequency across multiple days, consistent with positive selection (**Fig. 2C**), including the recurrent nonsynonymous L628M mutation. Other consensus changes did not persist within a host for more than one day with the exception of PA F9F and HA S95Y, which were present at two sequential time points (**Supp. Fig. 2A**). Interestingly, we observed similar allele frequency dynamics for the L628M and T109I mutations in a single ferret (**Fig 2C**). Surprisingly, the L628M and T109I mutations also co-occurred in 6 other ferrets in which they also tended to have similar, albeit lower, allele frequencies (**Fig 2D**).

**Fig 2.**
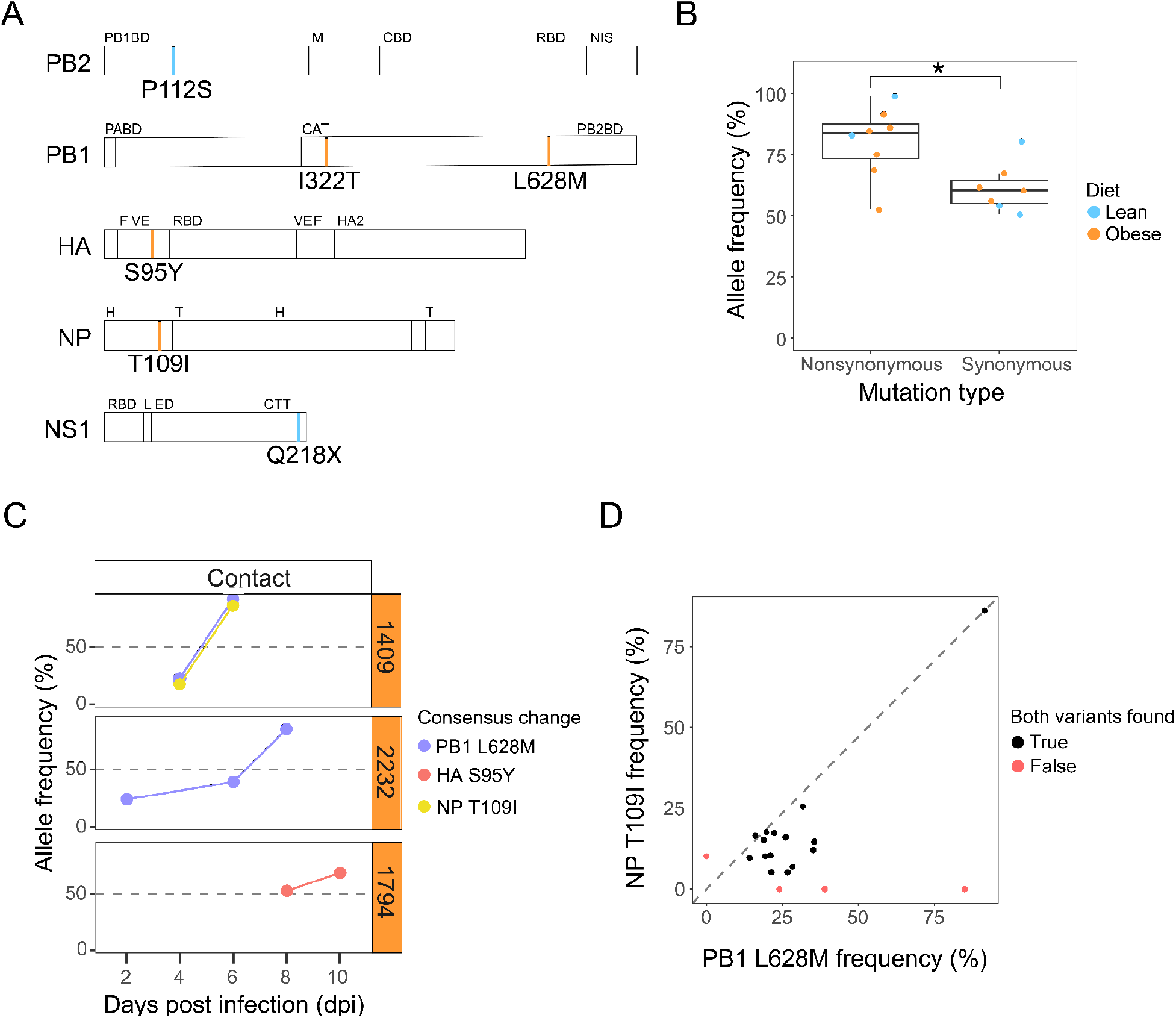
Properties of consensus sequence changes in infected ferrets. **A)** Schematic of protein domains containing nonsynonymous consensus changes. PB1BD = PB1 binding domain, M = middle, CBD = cap binding domain, RBD = receptor binding domain, NIS = nuclear import signal. PABD = PA binding domain, Cat = catalytic domain, PB2BD = PB2 binding domain. F = fusion domain, VE = vestigial esterase domain, HA2 = hemagglutinin 2 domain. H = head domain, T = tail domain. L = linker domain, ED = effector domain, CTT = C terminal tail. The predicted amino acid change of the consensus variants is labeled. The Q218X nonsense consensus change on the NS segment is specific to the NS1 splice form. It is also predicted to result in a synonymous (N60N) change in NS2, which is not included in the figure for clarity. **B)** Allele frequencies of nonsynonymous and synonymous consensus changes. Color indicates the diet of the ferret. * = p < 0.05 using a two-tailed Student’s t test. **C)** Allele frequencies of consensus changes that increased over sequential days within a ferret. **D)** Frequency of L628M and T109I mutations at each timepoint when they co-occurred (n = 15, black points) and when they did not co-occur (n = 4, red points) in a total of 6 ferrets.

### Obesity selects for unique minor variants

In addition to consensus sequence changes, there were a large number of minor variants (n = 567) that were not in the stock but arose *de novo* within a ferret (**Table S3**). Initially, we focused on minor variants that were shared between ferrets of the same diet group as these may be the result of diet-specific adaptation. Shared minor variants (n = 26) occurred throughout the genome (except MP) and were predominantly nonsynonymous (n = 19) (**Fig. 3**). Variants shared between multiple individuals can result from either recurrence or transmission. Most shared nonsynonymous minor variants were the result of recurrence as they were observed in unrelated ferrets (n = 14, purple points in **Fig. 3**). A small number of shared nonsynonymous minor variants were found both in an index ferret and the corresponding contact ferret, and are therefore either the result of independent recurrence or transmission (n = 5, green points in **Fig. 3**). All shared nonsynonymous minor variants appeared transiently as we identified only one case in which a variant was present in two sequential time points within a ferret **(Supp. Fig. 2B**).

**Figure 3.**
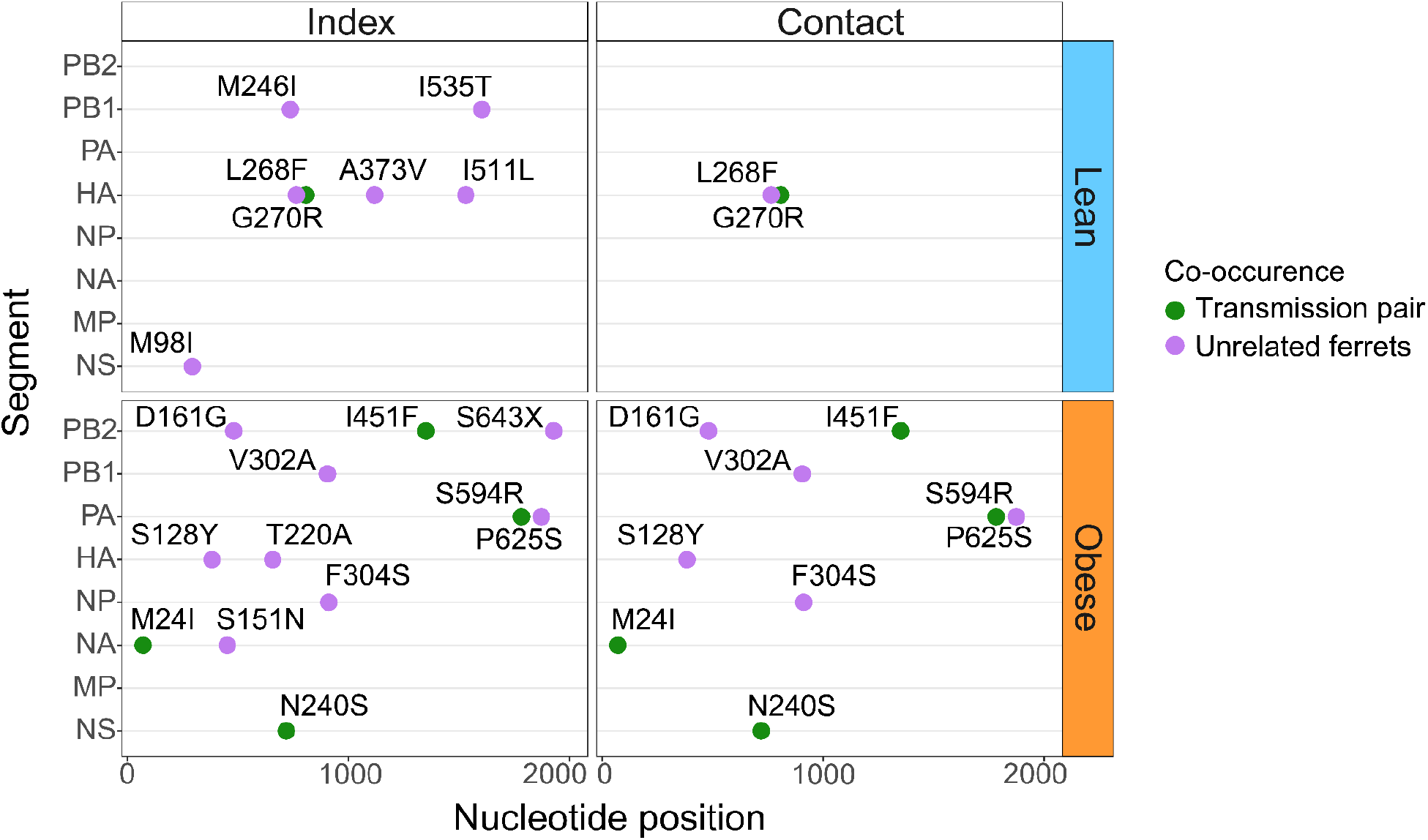
Recurrent and transmitted diet-specific nonsynonymous minor variants. The nucleotide position of diet-specific *de novo* minor variants found in two ferrets and the corresponding amino acid change are indicated. Color indicates whether the variant co-occurred in a transmission pair or in unrelated ferrets. Co-occurrence within a transmission pair could be the result of either transmission or recurrence whereas co-occurrence in unrelated ferrets is likely due solely to recurrence. Facets indicate the diet and infection route of the ferret.

### Dynamics of viral diversity over the course of infection

We next sought to quantify the dynamics of viral genetic diversity over the course of infection. Index ferrets were best suited for addressing this question as each individual was infected with similar initial diversity as a result of pre-existing genetic variation in the inoculum and infection with a large inoculum (10^6^ TCID_50_). By contrast, the diversity in contact ferrets depended on the variation in the index ferret and the extent to which it was transmitted, which varied greatly between individuals. In our experimental design, variants present in the inoculum were considered standing genetic diversity, whereas *de novo* variants that arose within a ferret reflected new genetic diversity.

We quantified viral genetic diversity using the consensus changes and minor variants in index ferrets (n = 12 obese, n = 11 lean) using several metrics. First we calculated SNV richness in each ferret at each time point (**Fig. 4A**). The richness at 2 dpi was similar to the inoculum and subsequently increased over time, peaking at 4 dpi in both lean and obese index ferrets as a result of the acquisition of *de novo* variants that were not present in the inoculum. Variants were subsequently lost as the infection was cleared and the total viral population decreased as reflected by the decrease in SNV richness detectable at 6 dpi. The overall dynamics were similar for nonsynonymous and synonymous variants and did not differ between diet groups. Comparisons of SNV richness for each segment individually also showed no significant differences between obese and lean ferrets (**Supp. Fig. 4A**). We also calculated diversity using Shannon entropy, which takes into account both the number of variants and their frequencies. This diversity metric showed similar dynamics to SNV richness with a peak at 4 dpi and no significant differences between diet groups (**Fig. 4B**).

**Figure 4.**
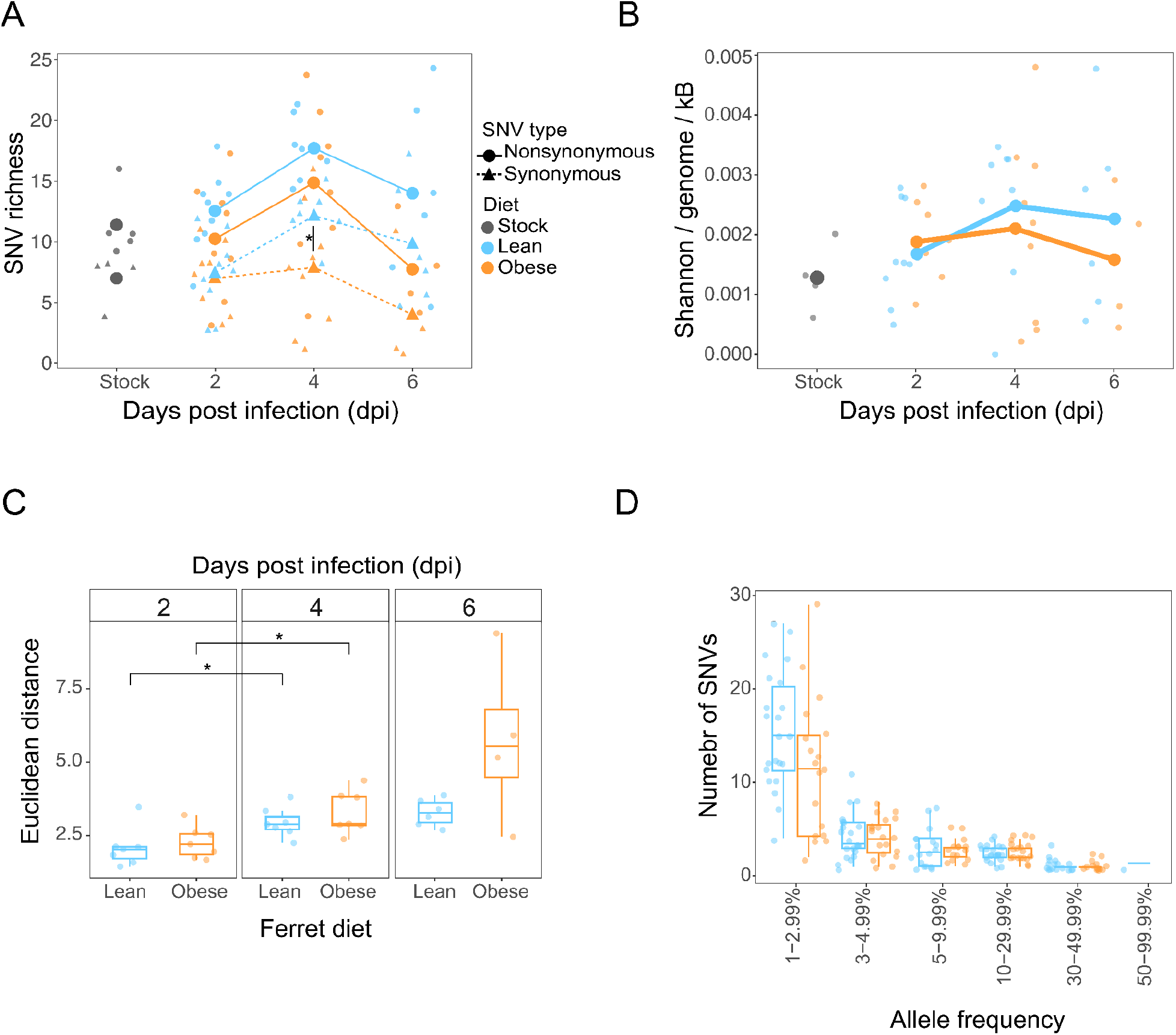
The dynamics of viral genetic diversity did not differ between lean and obese index ferrets. **A)** Richness per sample calculated by counting the number of SNVs in each index ferret sample. Small points indicate values for individual samples, while larger points connected by lines indicate average values for all samples within a diet group. Color indicates the diet of the ferrets. Dashed lines and triangle points indicate synonymous SNVs while solid lines and circle points indicate nonsynonymous SNVs. All comparisons were not significantly different using a two-tailed Student’s t test unless indicated * = p < 0.05. **B)** Shannon entropy per genome per kilobase calculated using the allele frequencies at each SNV position in each index ferret sample. Color indicates the diet of the ferret. **C)** L2-norm distance (Euclidean distance) calculated between each sample and its associated stock by summing the frequencies of all four nucleotides at all positions within a sample, reflecting all SNVs. Facets indicate dpi and color indicates the diet of the index ferrets. **D)** Allele frequency distributions of SNVs in index ferrets. Color indicates the diet of ferrets.

In contrast to SNV richness and Shannon diversity, the genetic distance between nasal wash samples and the inoculum continued to increase over the course of infection (**Fig. 4C**). This was a result of the simultaneous processes of purging standing genetic variation and introduction of new variation through *de novo* mutations. A substantial fraction of standing genetic variation was maintained in index ferrets over multiple sequential days of infection; however, their frequencies did not differ by more than 10% from their original frequency in the inoculum, suggesting an absence of strong directional selection acting on standing genetic variation (**Supp. Fig. 4B**). At the same time, 46% of variants that were present in the inoculum were not detected at the earliest time point in an index ferret, regardless of diet (**Supp. Fig. 4C**) and most variants were only detected at a single time point (**Supp. Fig. 4D**) indicating a remarkably rapid rate of turnover. The distribution of allele frequencies of these variants was highly skewed to low frequencies (less than 5%), consistent with a high rate of introduction of new mutations (**Fig. 5D**). Overall, we do not find that the dynamic changes in genetic diversity were impacted by diet.

**Figure 5.**
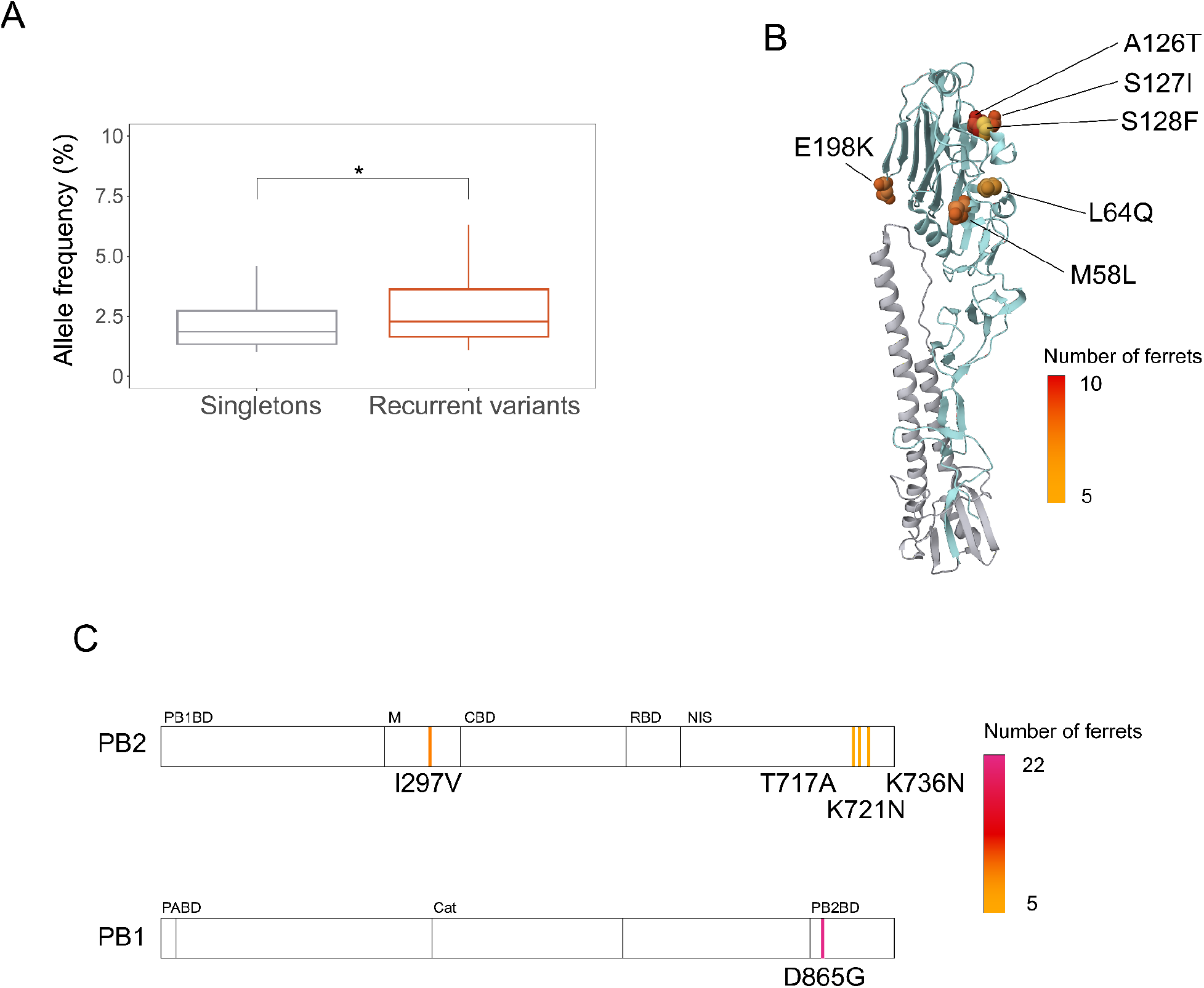
*De novo* nonsynonymous minor variants identified in multiple ferrets independent of diet. **A)** Allele frequencies of *de novo* minor variants that were present in both obese and lean ferret hosts compared with *de novo* variants that were present in only one ferret. * = p < 0.05 using a two-tailed Student’s t test. **B)** Location of shared *de novo* nonsynonymous minor variants mapped onto the 3D structure of the HA protein. Labels denote the amino acid change. Color represents the number of ferrets in which the variant was found. The HA structure of A/swine/Hong Kong/9/98 (H9N2) is shown in its cleaved form (Ha et al., 2001). **C)** Position of shared *de novo* nonsynonymous minor variants in PB2 and PB1 segments. Labels denote the corresponding amino acid change. Color represents the number of ferrets in which the variant was found. PB1BD = PB1 binding domain, M = middle, CBD = cap binding domain, RBD = RNA binding domain. NIS = nuclear import signal. PABD = PA binding domain. Cat = catalytic domain. PB2BD = PB2 binding domain.

### Highly recurrent variants indicate adaptation to the ferret host

Recurrent generation of *de novo* variation and its positive selection is rare and therefore difficult to detect. However, the large number of ferrets in our study and the use of an avian-like influenza virus in a mammalian system increased our power to detect these rare events. There were multiple instances in which *de novo* variants were shared between ferrets in different diet groups. These recurrent variants were found at higher allele frequencies than minor variants that appear only in one individual (**Fig. 5A**). We identified a cluster of 6 highly recurrent variants in the HA1 domain of the HA protein, which contains residues important for binding to host cells (**Fig. 5B**). Additionally, we find a set of highly recurrent variants in the PB2 and PB1 segments, which encode components of the viral polymerase (**Fig. 5C**). Of particular interest is the nonsynonymous SNV in the PB1 segment, resulting in an aspartic acid (D) to glycine (G) change at amino acid position 685, which was present at low frequency (1-5%) in 22 out of 37 ferrets. These highly recurrent *de novo* nonsynonymous variants that occur in both diet groups likely reflect adaptation to the ferret host, rather than host diet-specific adaptations.

### DVG diversity is maintained throughout infection

DVGs have been shown to influence virus-host interactions and to be associated with stronger host immune responses (Wang et al., 2020). As with SNVs, the diversity of DVGs in contact ferrets may be dependent on both host diet and variation transmitted from the index ferret. To investigate the effect of host diet alone on DVG diversity, we performed DVG detection in index ferret samples using the DiVRGE pipeline (see **Methods**). The vast majority of unique DVGs occurred in the three polymerase gene segments (**Fig. 6A**). Following the initial infection event in index ferrets, DVG richness did not change significantly over the course of infection, regardless of host diet. However, when considering only *de novo* DVGs that were not present in the inoculum, we detected multiple DVGs that arose specifically within a host diet group (**Fig. 6B**). These DVGs ranged in size from 473 nucleotides to 2034 nucleotides long – almost the entire length of the PB2 gene – and in general were symmetrical around the segment midpoint with conserved segment ends. There was no significant difference in the size distribution between DVGs that occurred in obese or lean ferrets for any segment. While *de novo* DVGs were typically found in a single ferret, we observed instances in which the same DVG was found recurrently within obese or within lean ferrets (**Fig. 6C**). The most highly shared DVGs within lean ferrets occurred in the PA segment, whereas the most highly shared DVGs within obese ferrets occurred in the PB2 and PB1 segments, possibly reflecting differential selection pressures as a result of host diet.

**Figure 6.**
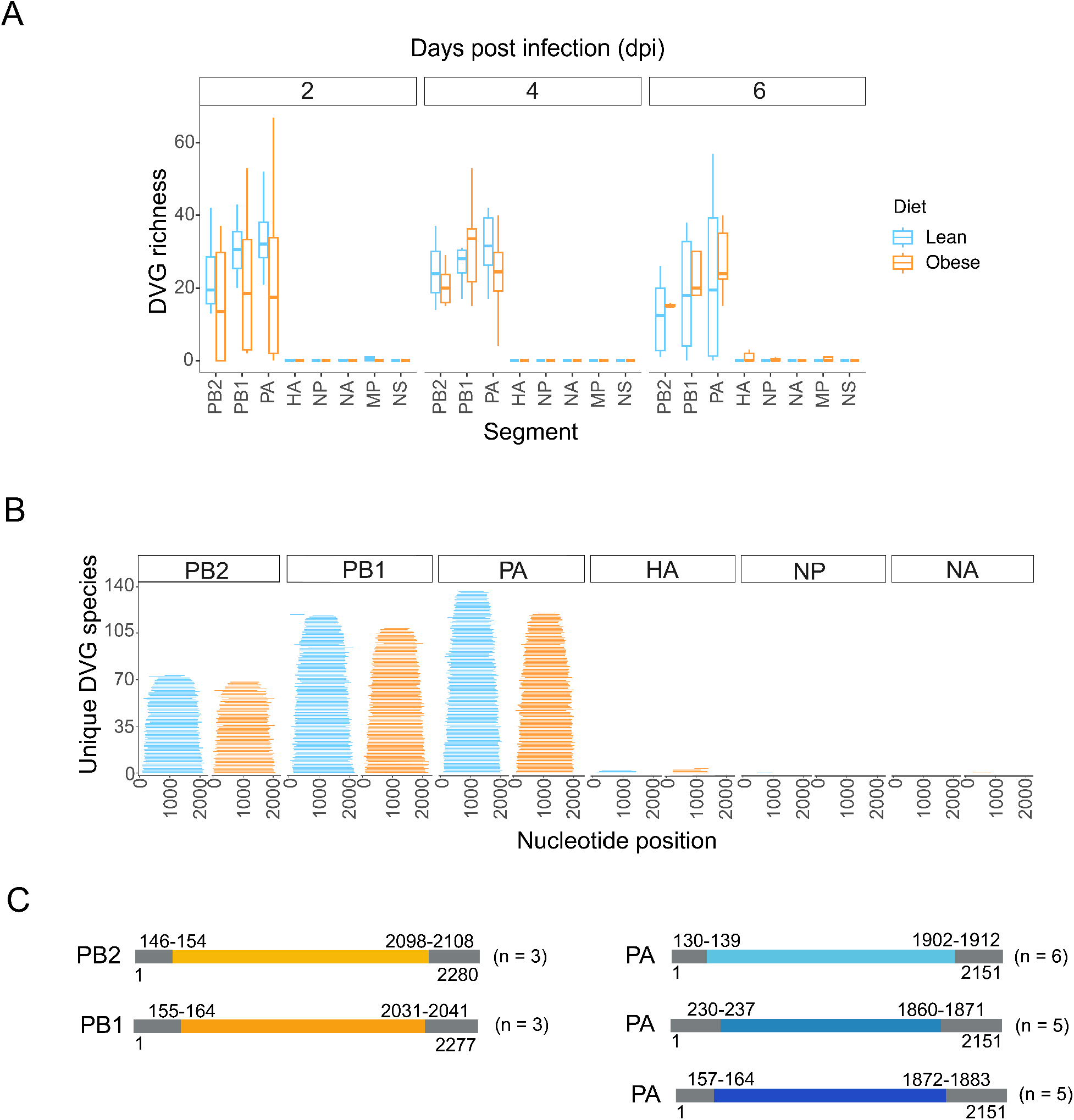
Diversity of defective viral genomes in index ferrets. **A)** DVG richness per sample calculated by counting the number of unique DVGs in each sample. Facets indicate dpi and color indicates diet of index ferret. **B)** Size distribution of the DVGs unique to either obese or lean ferrets. Each line represents the deleted region in a DVG, extending from the start to the end of the deletion. Color indicates diet. Facets indicate segment. **C)** The most common DVGs within either obese or lean ferrets, with the predicted deletion start and endpoints annotated. The coordinate range of breakpoints is a result of the DiVERGE analysis pipeline grouping DVGs with similar breakpoints. The number of individual ferrets (n) in which the DVG is found is indicated.

## Discussion

In this study we sought to understand the implications of host obesity on IAV intrahost evolution. Obesity is associated with significant immune system dysfunction and therefore the environmental conditions within the host in which the viral population evolves may be distinct from that of non-obese individuals. To address this question using a controlled experimental approach, we made use of an obese ferret model. We found evidence for specific adaptation of IAV to an obese host through both recurrent and high frequency variants that were unique to obese individuals. By contrast, we did not find that overall IAV genetic diversity differed between obese and lean hosts, highlighting the necessity of considering the specific mutations, their frequencies, and dynamics in the context of host factors. By using an avian-like strain of H9N2 in a ferret model we also observed signatures of host adaptation independent of metabolic state.

Our study design enabled the identification of a unique class of nonsynonymous variants in obese hosts that arose *de novo* within an individual, showed systematic increases in allele frequency over the course of infection, and ultimately became the dominant variant. Importantly, we identified this class of variant only in obese contact ferrets. The three specific mutations that exhibited this behavior, PB1 L628M, NP T109I, and HA S95Y, initially appeared as minor variants in the infected contact ferrets but were not transmitted from the index ferret. This suggests that transmission did not play a role in the selection of these variants, but rather that they arose *de novo*, and increased in frequency, within the contact ferret. These unique dynamics observed in obese contact ferrets are likely due to the differences between mechanisms of infection in our experiment. The index ferrets were intranasally inoculated with a very large, genetically diverse viral population. However, it is estimated that only 2-3 viral particles are passed from index to contact during a transmission event (Ghafari et al., 2020; McCrone et al., 2018). Thus, infection in the contact ferrets was seeded by a very small viral population with low genetic diversity, which subsequently expanded and accumulated more variants during the course of infection. The small effective population size may allow a single variant to rapidly increase in frequency more easily than in the index ferrets. We did not identify a similar class of variants in lean contact ferrets, perhaps reflecting reduced strength of selection in these hosts.

PB1 L628M is especially relevant when considering adaptation to the obese host environment as it is a nonsynonymous change that was present in several lean and obese ferrets, but only swept to high frequency in obese ferrets. Surprisingly, the PB1 L628M mutation almost always co-occurred with the NP T109I mutation, which may be indicative of an epistatic interaction between these variants (Lyons & Lauring, 2018; Nshogozabahizi et al., 2017). During virion production, the viral RNA genome is wrapped around NP proteins and packaged with viral polymerases, which bind to the ends of each of the genomic segments (Samji, 2009). It is possible that the PB1 L628M and NP T109I variants stabilize this complex in a cellular environment with increased inflammatory gene expression, as found in obesity.

There were many SNVs and DVGs that were unique to either obese or lean ferrets that may play a role in mediating adaptation depending on metabolic state of the host. A total of 12 nonsynonymous minor variants were shared between obese ferrets either as a result of transmission or recurrence. Two of these variants are particularly noteworthy, HA S128Y and T220A, as they are located in the HA1 sialic-acid binding domain and could possibly enhance binding to epithelial cells in the altered cellular state of obesity or allow the virus to avoid detection by antibodies. Our observation suggests one possible effect of obesity is to enable selection of minor variants, even in the highly variable HA gene segment, that would have been selectively purged in lean hosts by expanding the exploration of evolutionary space and thereby accelerating antigenic drift. In addition to SNVs, there were multiple DVGs that were found recurrently within a diet group. DVGs in the PA segment were especially common in the lean ferrets, with several occurring in half, or more, of the lean ferrets. By contrast, even the most highly shared DVGs in obese ferrets occurred in only 3 individuals (i.e. < 30%), and were present on the other polymerase genes (PB1 and PB2). As a DVG needs to infect the same cell as a replication-competent virus in order to be propagated (Meng et al., 2017; Samji, 2009), these results suggest that obesity may interfere with the formation and spread of specific common DVGs and result in less common transient DVGs, perhaps due to the altered cellular environment found in obese ferrets.

Although we find extensive evidence for host-specific adaptation, we did not observe differences in the overall dynamics of genetic diversity in obese and lean hosts. In all ferret hosts we observe remarkably rapid turnover of diversity as reflected by the ephemeral nature of many SNVs. In index ferrets, diversity peaked 4 days after infection and subsequently declined as the infection was cleared. We also identified several candidate variants that are likely to underlie adaptation to the ferret host regardless of metabolic state as they were present in both lean and obese individuals. Changes in the polymerase gene segments (PB2, PB1, and PA) and the HA segment are associated with viral host switching and we observed several highly recurrent variants in these genes. Of particular interest is the D685G variant located on the PB1 segment that was found at low frequencies in 22 out of 37 ferrets. Additionally, there were several, highly recurrent variants in the HA1 domain of the HA protein, which may allow more efficient binding to ferret epithelial cells in the respiratory tract, which have different sialic acid distributions than avian species.

Although H9N2 is not currently a seasonal subtype of IAV in humans, it infects avian species and has caused sporadic cases of human disease. As such, it remains a subtype of public health concern. H9N2 has intermediate rates of transmission in the ferret model, which enables assessment of susceptibility to infection through transmission (Meliopoulos et al., *unpublished*). However, the relatively small number of transmission events in our study limited our ability to quantify the diversity of transmitted variants between obese and lean individuals. Nonetheless, our study provides evidence that minor variants are likely transmitted, especially in conditions where direct contact occurs for a period of time, potentially leading to multiple transmission events. Future studies will be aimed at assessing the effect of obesity on transmission dynamics and bottleneck sizes. It would also be important to study the extent to which our findings generalize to other IAV subtypes and to extend sampling beyond upper respiratory tracts to study tissue tropism. Finally, integration of host gene expression states may provide mechanistic insights into the molecular basis of differential selective pressures in lean and obese hosts.

## Methods

### Ferret raising protocol

Influenza-naive and neutered male 6-week-old ferrets were obtained from Triple F Farms. Ferrets were randomly assigned to the obese or lean diet groups and fed the appropriate diet group for a total of 12 weeks (Meliopoulos et al., *unpublished*). Total weight, waist circumference, and skinfold fat measurements were made weekly. All animal experiments were approved by the St. Jude Children’s Research Hospital Institutional Biosafety Committee and Animal Care and Use Committee (protocol 513) and were in compliance with the Guide for the Care and Use of Laboratory Animals. These guidelines were established by the Institute of Laboratory Animal Resources and approved by the Governing Board of the US National Research Council.

### Infection experiments and sequencing protocol

A/Hong Kong/1073/1999 (H9N2) virus was propagated in embryonated chicken eggs as previously described (Brauer & Chen, 2015) and aliquots were stored until use in infection experiments. At this time, they were thawed and diluted in phosphate buffered saline (PBS) prior to intranasal inoculation of index ferrets with 10^6^ TCID_50_ virus at day 0. On 1 dpi, a naive contact ferret was introduced in the same cage as an index ferret. Ferrets remained co-caged through the end of the experiment at 12 dpi. Experiments were performed using five different cohorts and the results aggregated. Nasal wash samples were collected from anesthetized index and contact ferrets every second day starting at 2 dpi until 12 dpi. Sneezing was induced in anesthetized ferrets (30 mg/kg ketamine delivered intramuscularly, Patterson Veterinary Supply) by instillation of 1 mL PBS supplemented with antibiotics (100 U/mL penicillin and 100 ug/mL streptomycin) to the nasal cavity. Sample was collected, briefly centrifuged, and stored at −80°C. Viral RNA was extracted from 50 µL of nasal wash or the viral inoculum using the Qiagen QIAmp Viral RNA Mini Kit (QIAGEN, Cat#52904). Viral titers were determined by TCID_50_ analysis.

qPCR analysis was performed using primers targeted against the influenza matrix (M) gene segment: forward 5’-GACCRATCCTGTCACCTCTGAC-3’, reverse 5’-AGGGCATTYTGGACAAAKCGTCTA-3’, probe 5’-TGCAGTCCTCGCTCACTGGGCACG-3’ using the following conditions on a CFX96 Real Time PCR (Biorad): 50°C for 5 minutes, 95°C for 20 seconds, followed by 40 cycles of 95°C for 3 seconds and 60°C for 30 seconds. Copy number of the M gene segment was determined by comparison to a standard curve generated using a FluA gBlock fragment (5’-TCGCGCAGAGACTGGAAAGTGTCTTTGCAGGAAAGAACACAGATCTTGAGGCTCTCATGGAATGGCTAAAGACAAGACCAATCTTGTCACCTCTGACTAAGGGAATTTTAGGATTTGTGTTCACGCTCACCGTGCCCAGTGAGCGAGGACTGCAGCGTAGACGCTTTGTCCAAAATGCCCTAAATGGGAATGGGGACCCGAACAACATGGATAGAGCAGTTAAACTATACAAGAAGCTCAAAAGAGAAATAACGTTCCAT-3’). RNA concentration was measured using a spectrophotometer (Nanodrop).

Viral genomic RNA was amplified using universal Influenza A primers: Uni 12/Inf 1 5’-GGGGGGAGCAAAAGCAGG-3’, Uni12/Inf3 5’-GGGGGGAGCGAAAGCAGG-3’, and Uni13/Inf 1 5’-CGGGTTATTAGTAGAAACAAGG-3’ (IDT). Multiplex reverse transcription-PCR (M-RTPCR) was performed as previously described (Zhou et al. 2009). Amplification was confirmed with gel electrophoresis using a 1% agarose gel stained with ethidium bromide. Samples were diluted to a concentration of 0.2 ng/µL and sequencing libraries were prepared using the Nextera XT Library Prep protocol (Illumina FC-131-1024). M-RTPCR and library preparation was performed in duplicate for all viral RNA. Samples were pooled and sequenced on an Illumina NextSeq 500 using paired-end (2×150bp) mode (Illumina, Inc., San Diego, CA).

### Data processing

Sequence reads were trimmed using Trimmomatic v0.39 with the arguments LEADING:20 TRAILING:20 SLIDINGWINDOW:4:20 MINLEN:20 (Bolger et al., 2014) and aligned to the Influenza A/Hong Kong/1073/1999 (H9N2) reference genome using bwa-mem2 version 2.1 (Vasimuddin et al., 2019). Duplicate reads were marked using Picard v2.23.8 (http://broadinstitute.github.io/picard) and removed using Samtools version 1.11 (Li et al., 2009). SNVs were identified using an in-house variant calling pipeline, timo (https://github.com/GhedinLab/timo). Samples were required to have at least 200x coverage at 40% of positions in all eight genome segments. This ensured that samples with DVGs with low coverage in the middle of the segment were retained. In general, samples had segments with coverage in excess of 1,000x. 40% segment coverage was chosen so that all stock samples were marked as high quality, but samples were manually inspected to confirm their suitability for downstream analysis. Only variants present in both technical replicates were retained. Consensus changes were defined as SNVs at greater than or equal to 50% allele frequency and 10X coverage and minor variants were defined as those greater than 1% allele frequency (Roder et al., 2023). Defective viral genomes were identified using DiVRGE (https://github.com/GhedinSGS/DiVRGE) using a minimum deletion size of 5 nucleotides. Downstream analyses were performed in R. All analysis code can be found on GitHub: https://github.com/GreshamLab/ferret-host-obesity. Sequence data has been deposited in the SRA and is available at BioProject ID PRJNA993240.

### Diversity Calculations

Richness was calculated by summing the number of SNVs, regardless of frequency, present in each individual ferret at every time point for which sequencing data was acquired. We calculated richness with respect to the entire genome and each segment.

Shannon entropy was calculated using every SNV, regardless of frequency, for every individual ferret and time point. These values were summed for each sample, divided by the genome size, and normalized to kB by dividing the value by 1000.

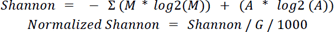

M: frequency of major allele. A: frequency of minor allele. G: genome size.

Euclidean distance (also known as L2 norm) was calculated using the dist() function in R. Briefly, the pairwise distance between two samples was determined using the frequency of every nucleotide summed across every position across the genome. Distance was calculated for all pairwise comparisons of samples.

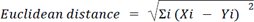

Xi: Allele frequencies across the genome of sample X. Yi: Allele frequencies across the genome of sample Y.

## Acknowledgements

We thank members of the Schultz-Cherry laboratory for their assistance with ferret experiments including Brandi Livingston, Bridgett Sharp, Sean Cherry, Lauren Lazure, and Dr. Ginna Hargest. This work was supported in part by the Division of Intramural Research (DIR) of the NIAID/NIH (EG) as well as the National Institutes of Health Centers for Excellence in Influenza Research and Surveillance (CEIRS) contract HHSN27220140006C (SSC), National Institutes of Health Centers for Excellence in Influenza Research and Response (CEIRR) contract 75N93021C00016 (SSC), the Center for Vaccine Research in High Risk Populations (CIVR-HRP) NIAID contract 75N93019C00052 (SSC and DG), National Institutes of Health grant R01AI140766 (SSC and DG), ALSAC (SSC), National Institutes of Health grant R01GM134066 (DG), and National Institutes of Health grant T32GM132037 (MK).

## Supplemental Figures

**Supplementary Figure 1.**
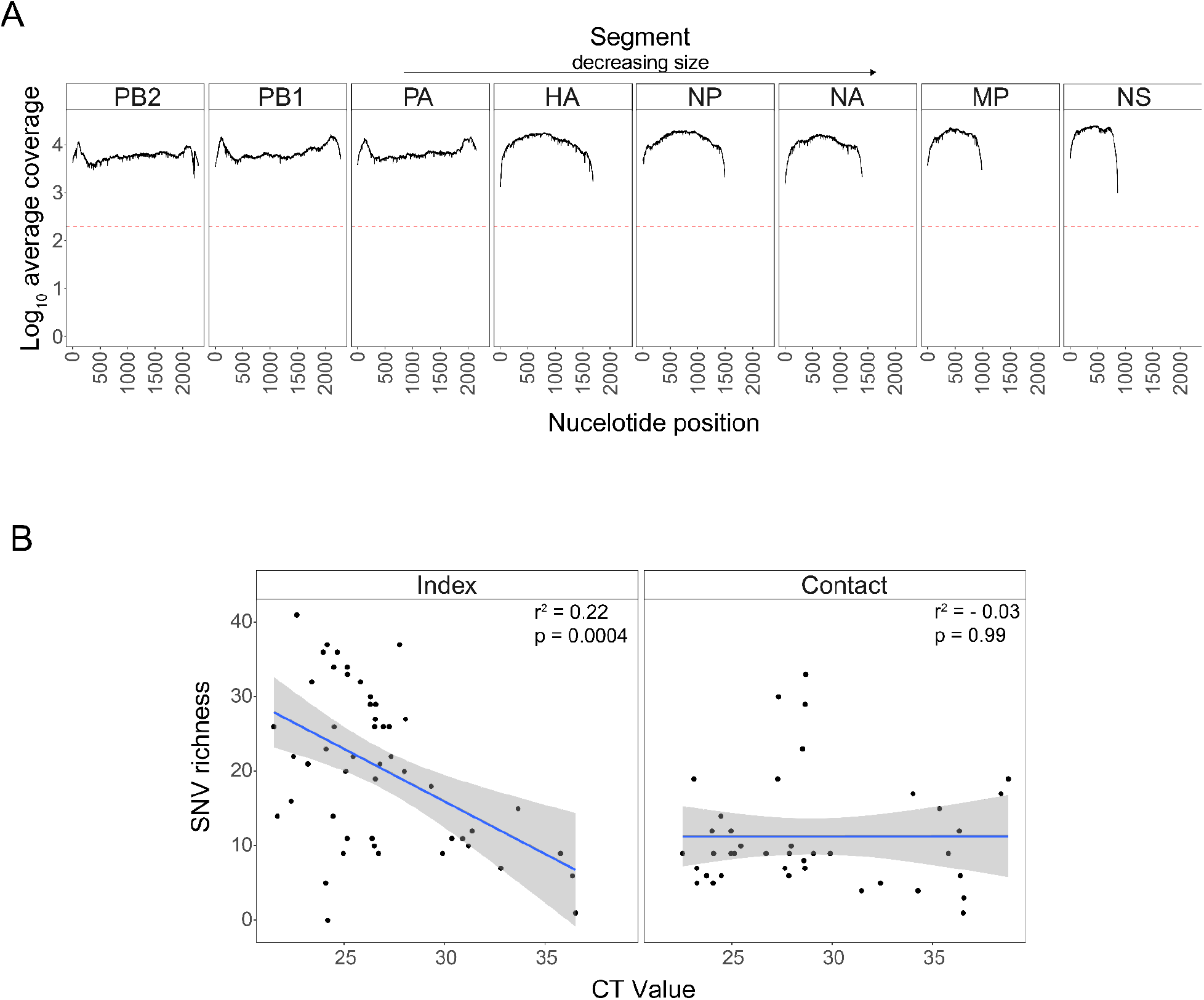
Quality control of viral sequencing data. **A)** Average coverage across each nucleotide position for all 8 segments of the IAV genome for all samples. Facets indicate segments, which are ordered in decreasing size. **B)** Correlation between qPCR Ct values and number of SNVs for each sample. A linear model was fit to the data and the Pearson correlation (r^2^) and associated p-value determined.

**Supplementary Figure 2.**
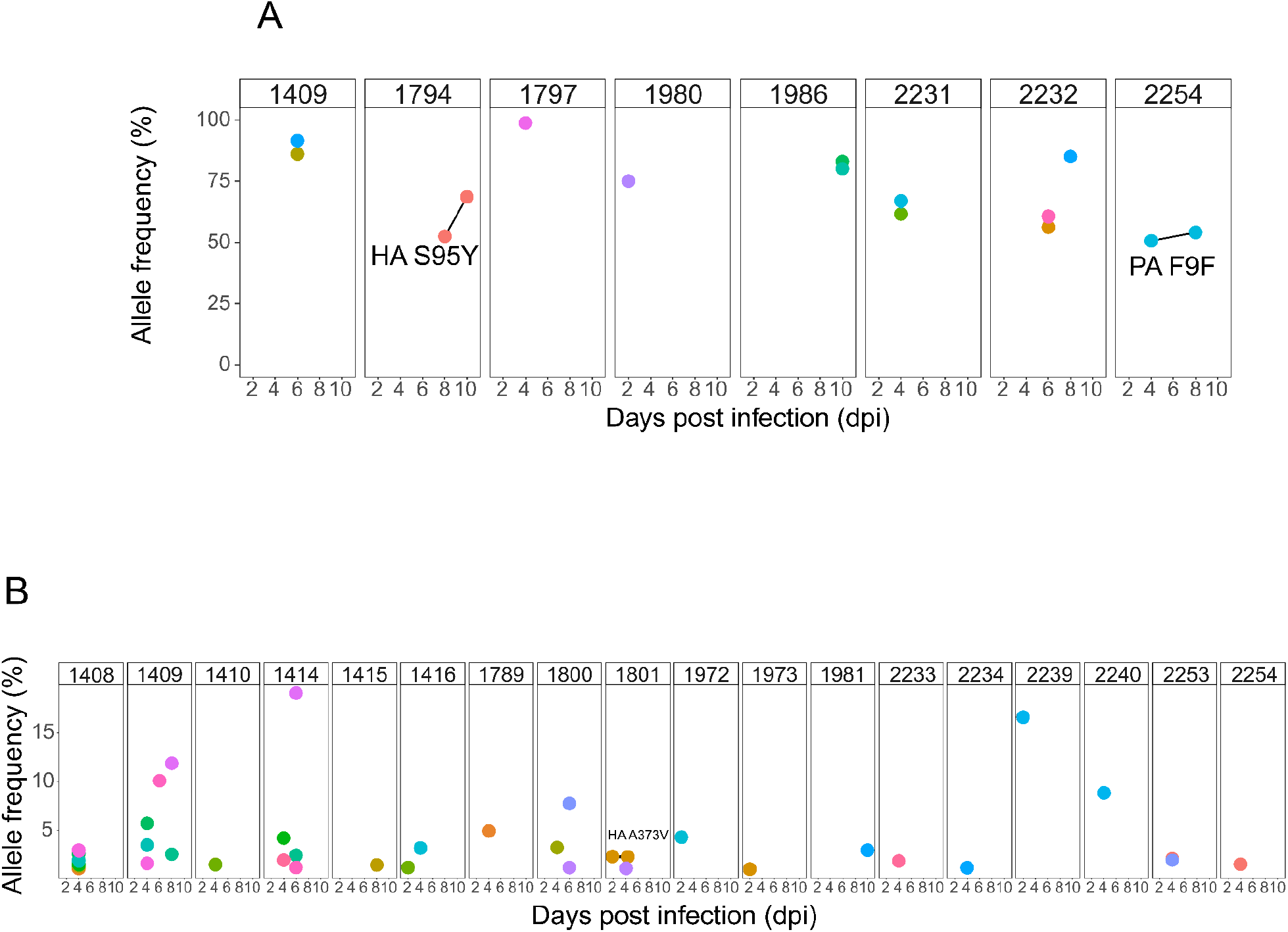
Persistence of consensus changes and minor variants. **A)** Allele frequency of all consensus changes. Lines connecting points indicate persistence of a variant for at least two time points. Persistent variants are labeled with their amino acid change. **B)** Allele frequency of recurrent *de novo* nonsynonymous minor variants identified in obese ferrets. Line connecting points indicate persistence of a variant for at least two time points. Persistent variants are labeled with their amino acid change.

**Supplementary Figure 3.**
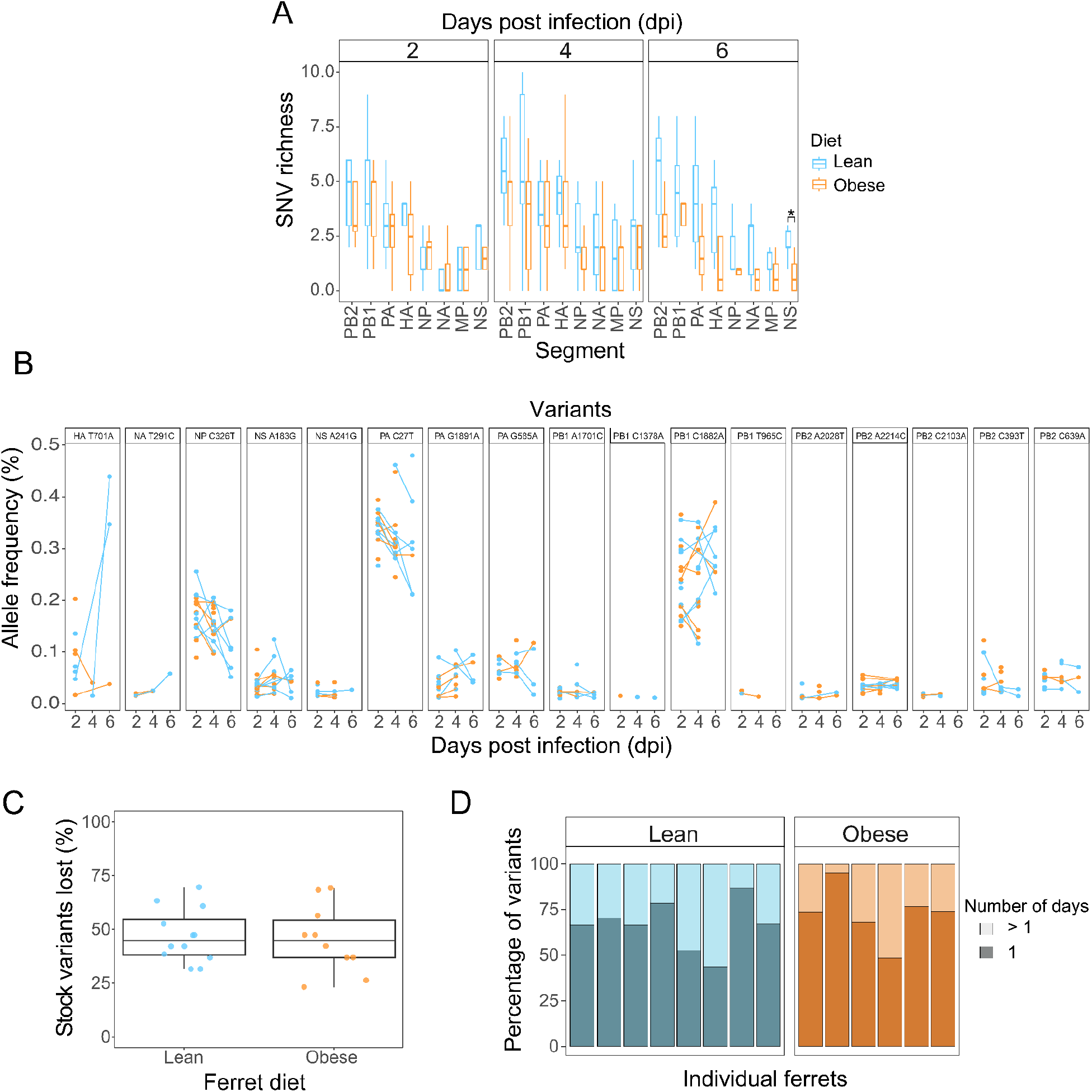
SNV dynamics in index ferrets. **A)** Distribution of SNV richness for each genome segment over all samples for days 2-6 dpi. **B)** Turnover of minor variants in index ferrets. Facets and color indicate diet of index ferret, shading indicates whether a variant was found in only a single time point (dark) or multiple time points (light) for the same ferret. **C)** Percentage of minor variants present in the stock that was identified in each index ferret. **D)** Variants that were identified at only one time point or at multiple time points.

## Supplementary Tables

**Table S1.** Metadata for all ferrets included in the study with physiological measurements taken prior to infection.

**Table S2.** Transmission pair partners.

**Table S3.** SNVs identified in each ferret, their amino acid change, and frequency.

## Supplementary Files

**Supplementary File 1.** Final output of the timo variant calling pipeline used for all downstream analyses performed in R.

**Supplementary File 2.** Output of the timo variant calling pipeline containing all positions in all samples at which any sample contained a variant for the purpose of calculating genetic distances.

